# Complete assembly of novel environmental bacterial genomes by MinION^TM^ sequencing

**DOI:** 10.1101/026930

**Authors:** Daniel J Turner, Xiaoguang Dai, Simon C Mayes, Sissel Juul

**Affiliations:** Oxford Nanopore Technologies Ltd., Oxford, UK; Oxford Nanopore Technologies Inc., New York, USA

## Abstract

In this study, we adapt a protocol for the growth of previously uncultured environmental bacterial isolates, to make it compatible with whole genome sequencing. We demonstrate that in combination with the MinION sequencing device, complete assemblies can be derived, allowing genomic comparisons to be made. This approach allows rapid, inexpensive and straightforward discovery, and genomic analysis, of previously uncultured prokaryotic genomes, and brings greater ownership of all parts of the sequencing process back to individual researchers.

---

There are an estimated 10^5^ to 10^6^ microbial species in existence, but only a few thousand such species have been isolated and cultured, because of the difficulty of growing many microbes under laboratory conditions. One reason for this difficulty is thought to be that many microbial species have specific nutritional requirements, without which they cannot grow ^1^. This is unfortunate because uncultured microbial species: i) have potential uses in biotechnology and in the development of new antibiotics ^2^, ii) contribute significantly to the biodiversity of all ecosystems, and iii) could be pathogenic, and hence warrant further study. Comparison of bacterial genome sequences helps us to understand genome evolution and gene function ^3^, and there is consequently a need for a quick, inexpensive and straightforward way to sequence and assemble uncultured prokaryotic genomes.

One way to derive genome sequences from complex samples is by metagenomic assembly, where total DNA is extracted from a sample and sequenced, and the multiple genomes which constitute the sample are assembled together. However, due to the potentially large number of genomes present in such a sample, this approach can make less distinct the whole concept of what constitutes an assembled genome ^4^. In contrast to this comprehensive approach, the iChip ^5^ allows individual, previously unculturable, bacteria to be isolated and grown clonally, by exposing the cells to the environment from which they were taken, in such a way that nutrients can diffuse in, with the bacterial cells immobilised in gel. Following their initial isolation and growth, it has been reported that many previously unculturable bacteria can adapt to being cultured under laboratory conditions ^5^.

The iChip device has previously been shown to have utility for the identification of unculturable bacterial species by 16S sequencing ^1^ and the discovery of new antibiotics ^5^. We sought to exploit the device to culture, for whole genome sequencing, bacteria present in a stream close to the Oxford Nanopore office, Oxford, UK, at the confluence of the Littlemore Brook and the effluent from a sewage treatment facility, which we reasoned would contain a high bacterial count. After two weeks of growth in the iChip, submerged in the stream, we found that we were able to continue growth of the bacteria in low salt culture medium, which had been prepared with sterilised and filtered water from the stream, and also on petri dishes of low salt LB-agar, prepared in the same way. Using this approach we were able to grow, and extract DNA from, individual bacterial strains with the expectation that we would be able to sequence and ultimately assemble these genomes.

Unlike metagenomic assembly, complete assembly of single genomes is relatively straightforward. The criteria for a good quality assembly are well-defined: large contigs with as few misassemblies as possible ^4^. Long sequence reads have been shown to facilitate assembly of long contigs, and even complete single contig assemblies, because they span repetitive elements in genomes ^6^.

The Oxford Nanopore MinION is a portable, inexpensive single molecule nucleic acid sequencing device, which plugs directly into the USB drive of a standard laptop and requires no additional infrastructure, and which is capable of generating very long reads in real time ^7^. Using the Celera assembler ^8^, preceded by one or more rounds of error correction, reads from the *Escherichia coli* strain K-12 obtained from the device can be assembled into a single contig assembly ^9,10^. We therefore applied this technology to the sequencing of several of the unculturable bacterial strains.

Initially, we attempted to identify the bacterial species present in the different isolates using Kraken ^11^, in real time (i.e. as reads were being generated by the MinION, **Table 1**. However, the results were fairly inconclusive, indicating that the majority of these particular bacterial subspecies are not present in the NCBI database. In a similar way to digital PCR ^12^, which is quantitative because single template strands are amplified in separate wells or droplets, it is evident from the number of times certain species were detected (**Table 1**.) that those bacterial strains were more abundant in the original sample than others. Here, *Pseudomonas* species were the most abundant.

**Table 1.**
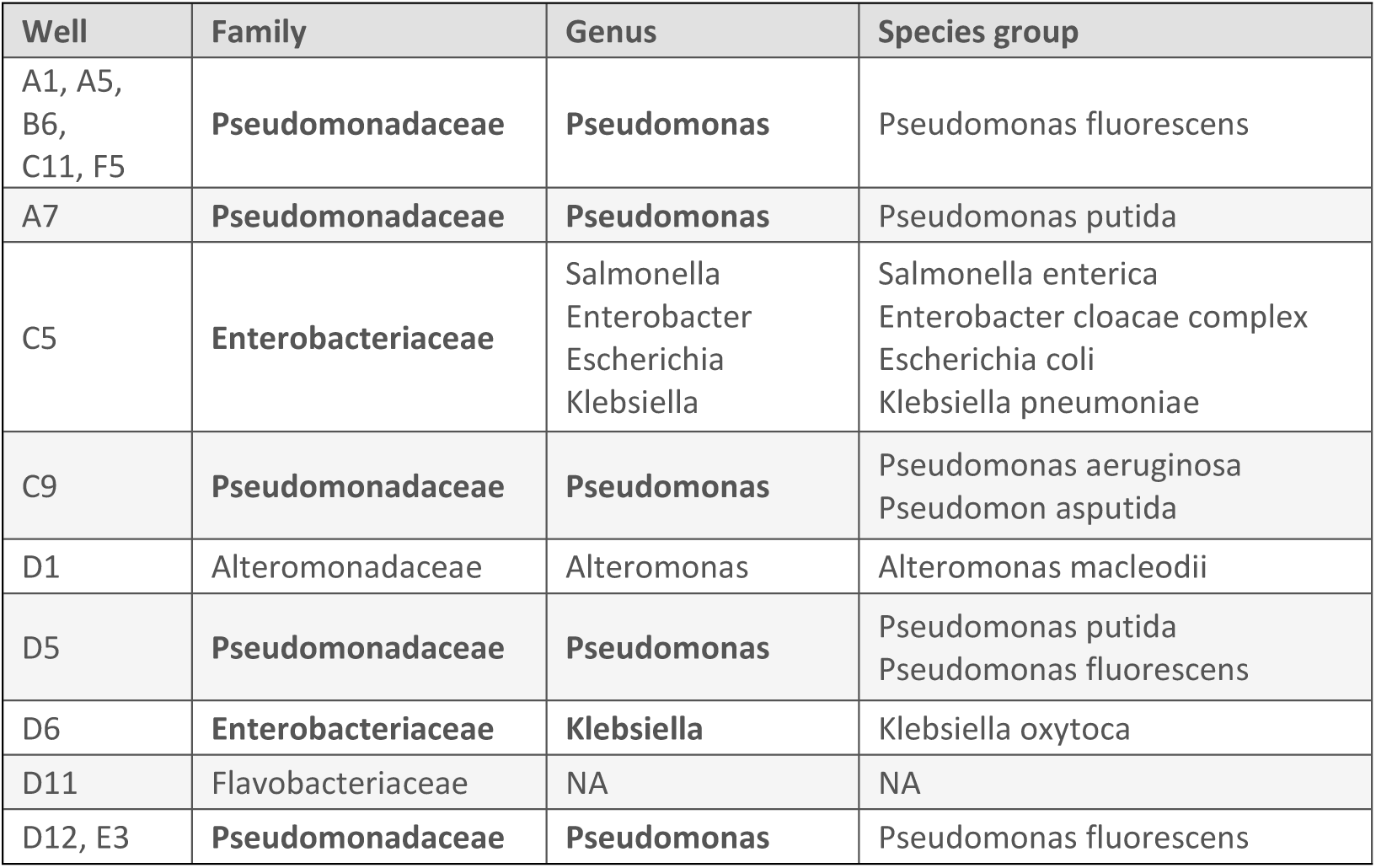
Bacterial isolates identified in the stream water using Kraken

We chose two isolates at random, C9 and D6 (**Table 1**.), which we had identified as strains of *Pseudomonas* and *Klebsiella oxytoca* respectively. We sequenced these genomes using PCR-free libraries with fragment lengths of approximately 8-10 kb, and we generated around 180Mb of data for each, equivalent to around 30x coverage of the *K. oxytoca* genome. After sequencing, we performed error correction using Nanocorrect ^9^, assembly using the Celera assembler ^8^, and manual finishing. We obtained a circular contig from each strain, covering approximately 4.4 Mb for C9 (N50=4.4Mb) and 6 Mb (N50 = 5.7Mb).

We then performed whole genome BLAST ^13^ to find out where these strains lie in the taxonomy. By this measure the closest match of D6 was to the strain KONIH1 (**Fig 1a**.). A Mummer plot ^3^ of the assembled genome against the KONIH1 strain reveals that the two strains are different, but highly similar (**Fig 1b**.). Whole genome BLAST of C9 revealed some similarity to *Pseudomonas alcaligenes NBRC* 14159, though a fully assembled version of this genome is not currently available for further comparison. A mummer plot of C9 against *Pseudomonas pseudoalcaligenes* CECT 5344 indicated significant differences (**Supplementary Figure 1**). We performed BLAST of the 16S gene of C9, and obtained many hits to various *Pseudomonas* species (**Supplementary Table 1**).

**Figure 1.**
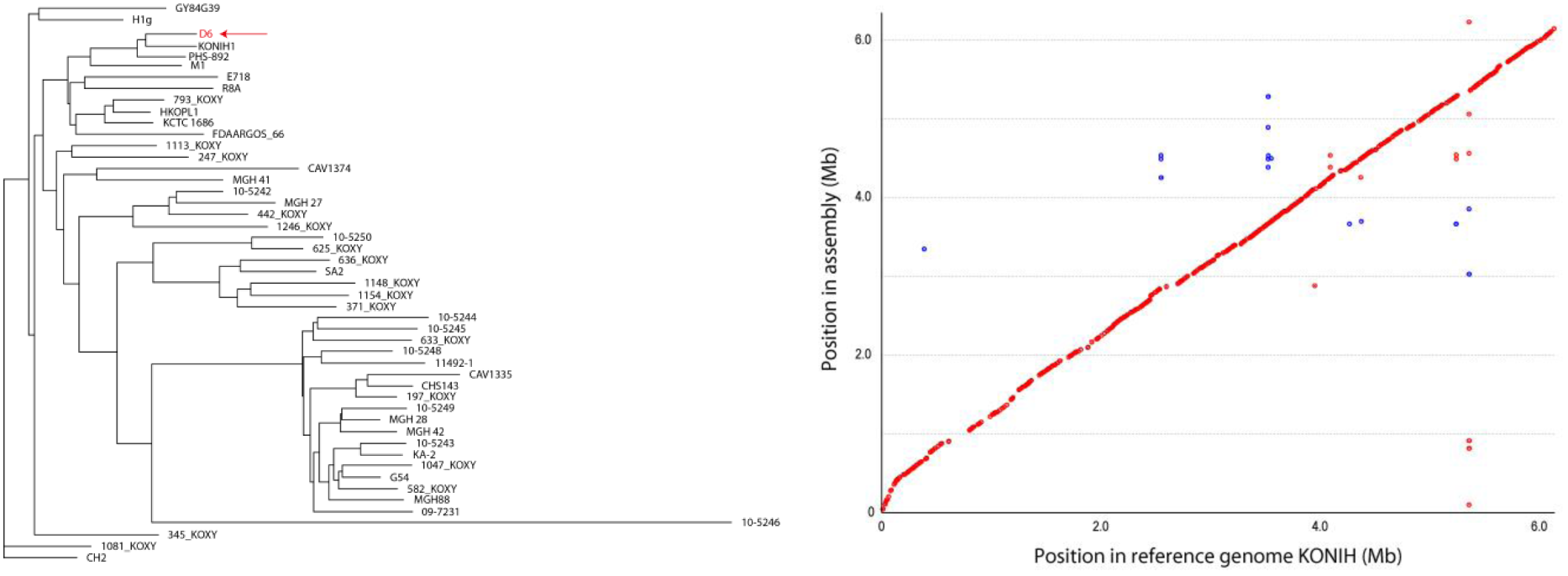
Genome comparison of one assembled bacterial strain to reference strains. **a)** taxonomy tree obtained by whole genome BLAST, with the assembled genome shown in red, **b)** Mummer plot of the assembled genome against the most closely matching reference genome.

In this study, we have adapted the iChip protocol to allow larger-scale growth of individual bacterial isolates, making it possible to obtain sufficient DNA for whole genome sequencing, and we have demonstrated that, in combination with Oxford Nanopore’s MinION, whole genome assemblies can be derived, allowing genome comparisons to be made. This will also enable further study of these bacterial strains, which had not previously been isolated or sequenced. The MinION device itself brings genome sequencing within the reach of small laboratories, and brings greater ownership of all parts of the sequencing process back to individual researchers. The combination of the iChip and the MinION will allow rapid, inexpensive and straightforward discovery and genomic analysis of previously uncultured prokaryotic genomes. It is important to note that the slowest step in this procedure is currently the culturing of the ‘unculturable’ bacteria: DNA extraction and library preparation can be performed in a morning, and sequence information is generated by the MinION in real time, starting virtually as soon as the first strand of DNA has translocated through a nanopore. Consequently, removal of the need to culture the bacteria could bring the time-to-result as low as a few hours. As throughput increases, the long read capability of the MinION will make metagenomic approaches the quickest route to assembling and characterising unculturable genomes.

## Method

We made an acrylic iChip ^5^ by 3D printing (Projet HD), dipped the central panel into 1% low melting point agarose (Invitrogen), allowed the gel to set at room temperature, and removed excess agarose with a scalpel. We then took a sample of water from the stream behind the Oxford Nanopore offices in Oxford, UK., dipped the iChip panel into the water sample, and assembled the device as described ^5^. We left the assembled iChip for 14 days in the stream, after which, we cleaned it externally and dismantled it. Using sterile cocktail sticks, we pushed agarose plugs out of iChip wells into separate wells of a deep-well 96-well plate (Bio-Rad) containing low salt LB medium (Sigma) which we had made with water from the stream, filtered (Corning) and sterilised by autoclaving. We sealed the plate with an air-permeable seal (Thermo Scientific) and left it on a plate shaker (Grant Bio) at 300rpm on a lab bench for two days so that the cultures would grow. We prepared Petri dishes containing low salt LB medium, again using filtered and sterilized stream water, but with 15 g/L agar (Sigma). We streaked out the cultures and allowed them to grow inverted on a laboratory bench for 2 days. Following this, we picked single colonies and grew them in 25ml liquid medium as previously described. We extracted DNA using a Qiagen 500Tip, following the manufacturer’s protocol, prepared PCR-free sequencing libraries using Oxford Nanopore Technologies’ version 6 library preparation kits, following Oxford Nanopore’s protocols, and sequenced each genome on separate flow cells following Oxford Nanopore’s standard running and analysis protocols.

We error-corrected data from each genome twice using NanoCorrect ^9^, and assembled the corrected reads using PBcR ^8^. We performed whole genome BLAST using discontiguous megablast ^14^, and generated genome alignment plots using Mummer ^3^

## Acknowledgements

We are grateful to Ameet Pinto for the stl files used for 3D iChip printing, and to Anthony Jones for printing the iChip.

